# Evaluation and optimization of the protocols for measuring cytochrome P450 activity in aphids

**DOI:** 10.1101/299545

**Authors:** He-He Cao, Tong-Xian Liu

**Affiliations:** Key Laboratory of Northwest Loess Plateau Crop Pest Management of Ministry of Agriculture, College of Plant Protection, Northwest A&F University, Yangling, Shaanxi, 712100, China; College of Agronomy, Northwest A&F University, Yangling, Shaanxi, 712100, China; Vertebrate Pest Division, Bangladesh Agricultural Research Institute, Joydebpur, Gazipur-1701, Bangladesh

**Keywords:** *Acyrthosiphon pisum*, cytochrome P450, detoxification enzymes, 7-ethoxycoumarin, *Myzus persicae*, *Rhopalosiphum padi*

## Abstract

Cytochrome P450 enzymes play major roles in insect detoxification of plant toxins and insecticides. However, measuring P450 activity in aphids has variable success, and a reliable method is not available yet. In this study, we evaluated and optimized the method for measuring P450 activity in aphids using the 7-ethoxycoumarin as the substrate. First, we found that nicotinamide adenine dinucleotide phosphate and protective agents are not needed in the aphid P450 activity assay, and homogenizing the green peach aphid, *Myzus persicae*, in the microplate resulted in significantly higher P450 activities than those in Eppendorf tube. Homogenizing aphids in Eppendorf tube could grind tissues thoroughly and released uncharacterized compounds that could inhibit aphid and pig liver P450 activities, whereas aphids in the microplate likely could not be thoroughly ground and thus released fewer such inhibitors. Then, the microplate homogenization method was optimized to follows: one or two aphids were put into one well of the microplate and ground in phosphate buffer using pipette tips for 20 cycles, followed by addition of 7-ethoxycoumarin, and then incubated for 1 h at room temperature, after which glycine buffer-ethanol mixture was added to stop the reaction. This method is also suitable for the pea aphid, *Acyrthosiphon pisum*, and the bird cherry □oat aphid, *Rhopalosiphum padi.* These results emphasize the importance of considering inhibitory effect of endogenous compounds in insects on their P450 activity and provide one possible method to reduce this inhibitory effect.

## Introduction

Cytochrome P450 enzymes can be found in almost all living organisms. They play important roles in the metabolism of endogenous compounds such as steroids, vitamins, hormones, carbohydrates and fatty acids, and they are also heavily involved in detoxifying exogenous compounds such as drugs, plant toxins, and mutagen (Liu et al., 2015). The P450 gene expression and activity in insects can be induced by plant toxins and further may confer insect resistance to insecticides (Liu et al., 2015). For example, overexpression of P450 gene *CYP6CY3* was essential for the green peach aphid, *Myzus persicae*, in detoxifying nicotine in tobacco as well as neonicotinoid insecticides (Puinean et al., 2010). The elevated P450 activity in insects has been used as a biomarker for certain insecticide resistance (Liu et al., 2015). Therefore, it is important to accurately measure the activities of P450 enzymes in insects.

Some previous studies failed to measure P450 activity in aphids as well as in some other small insects (Philippou et al., 2010; Gottardi et al., 2016), probably because the P450 activity was too low to detect by traditional method. Another possibility is that some uncharacterized compounds, released during homogenization, might inhibit the P450 activity (Orrenius et al., 1971; Wilson & Hodgson, 1972; Gilbert & Wilkinson, 1975; Valles & Yu, 1996). In addition, the P450 activity of mites could only be detected in supernatant after low-speed centrifugation, suggesting that the general P450 activity assays carried out with microsomal fraction prepared from high-speed centrifuged tissue homogenate may not be suitable for all organisms (Pasay et al., 2009). Until now, no universal method for measuring P450 activity in aphids has been established, although in certain aphid species such measurement has been reported (Castaneda et al., 2009).

Most assays measuring P450 activity are based on the transformation of non- or low-fluorescent P450 substrates to highly fluorescent metabolites catalyzed by P450 enzymes (Desousa et al., 1995; Gottardi et al., 2016). Among these substrates, 7-ethoxycoumarin (7-EC, 7-Ethoxy-1-benzopyran-2-one) is the most widely used chemical for measuring P450 dependent 7-ethoxycoumarin-O-dealkylation (ECOD) activity (Gottardi et al., 2016). This substrate has been widely used for measuring P450 activities in insects, fish, mammals, and nematodes, although other substrates may also work (Castaneda et al., 2009; Pasay et al., 2009; Gottardi et al., 2016). Therefore, we chose 7-ethoxycoumarin as the substrate for measuring aphid P450 activity in this study.

The ECOD activity in *M. persicae* was difficult to measure because of endogenous inhibitors presented in *M. persicae* homogenate, which could even significantly reduce ECOD activity of rabbit liver homogenate (Philippou et al., 2010). We obtained a much higher ECOD activity by homogenizing *M. persicae* in the 96-well microplate than in the Eppendorf (EP) tube. Furthermore, we found that homogenizing *M. persicae* in the EP tube inhibited ECOD activity of pig liver, whereas no inhibitory effect was found when aphids were homogenized in the microplate. Accordingly, we evaluated and optimized the method measuring ECOD activity in *M. persicae* by homogenizing aphids in the microplate. The results suggest that homogenizing aphids in the microplate can significantly reduce the release of inhibitors and is suitable for measuring P450 activity in *M. persicae* as well as some other aphid species.

## Materials and methods

### Chemicals

Potassium dihydrogen phosphate (KH_2_PO_4_, CAS: 7778-77-0, purity: ≥ 99.5%), dipotassium hydrogen phosphate trihydrate (K_2_HPO_4_3H_2_O, CAS: 16788-57-1, purity: ≥ 99.0%), glycerol (CAS: 56-81-5, purity: ≥ 99.0%), and ethanol (CAS: 64-17-5, purity: ≥ 99.0%) were obtained from Guangzhou Jinhuada Chemical Reagent Co., LTD (China). Phenylmethanesulfonyl fluoride (PMSF, CAS: 329-98-6, purity: ≥ 99.0%) was aquired from Beijing Solarbio Life Sciences Co., LTD (China). DL-dithiothreitol (DTT, CAS: 3483-12-3, purity: ≥ 99.0%), ethylenediaminetetraacetic acid disodium salt dihydrate (EDTA, CAS: 6381-92-6, purity: ≥ 99.0%), and β-nicotinamide adenine dinucleotide phosphate reduced tetra (cyclohexylammonium) salt (NADPH, CAS: 100929-71-3, purity: ≥ 98%) were from Sigma-Aldrich (China). 7-Ethoxycoumarin (7-EC, CAS: 31005-02-4, purity: ≥ 99.0%) were bought from J&K Scientific Ltd. (China) and 7-hydroxycoumarin (7-OH, CAS: 93-35-6, purity: ≥ 98%) were from Shanghai Macklin Biochemical Co., Ltd. (China).

### Aphids

*Myzus persicae* was reared on cabbage plants *(Brassica oleracea* L. var. *capitata*, var. “Qingan 70”) under a 14:10 h L/D cycle at 22 ± 2°C and 50% relative humidity. The pea aphid, *Acyrthosiphon pisum*, was reared on broad bean and the bird cherry-oat aphid, *Rhopalosiphum padi*, was reared on wheat under the same conditions.

### Comparison of homogenization methods

To measure P450 activity, most studies homogenize insect tissues in a mortar or an EP tube with fitted pestles, which could grind tissues thoroughly. Another method is putting intact insects or their dissected tissues directly into phosphate buffer in a multi-well microplate and then break up with pipette tips or proceed with no further homogenization (Desousa et al., 1995; Gottardi et al., 2016). First, we compared these two homogenization methods, i.e. homogenizing aphids in an EP tube with fitted pestles, and in a 96-well microplate with pipette tips. For EP tube homogenization method, adult aphids were placed into a 1.5 mL EP tube and homogenized with a plastic pestle in ice cold 100 mM phosphate buffer (pH 7.5) containing protectants (1mM EDTA, 1 mM DTT, 1 mM PMSF and 20% glycerol). The homogenate was used for measuring P450 activity without centrifugation. Then, 2.5 mg equivalent aphid homogenates were added to each microplate well that contained 100 μL 7-EC solution (0.5 mM 7-EC dissolved in 100 mM phosphate buffer pH 7.5 containing protectants). For the microplate homogenization method, 2.5 mg adult aphids were placed into each well of the microplate and homogenized in 100 μL 100 mM phosphate buffer (pH 7.5) containing protectants using a 200 μL pipette tip. Then, 100 μL 7-EC solutions (0.5 mM 7-EC dissolved in 100 mM phosphate buffer pH 7.5) were added to each microplate.

After the mix of aphid homogenates with 7-EC, the reaction was initiated by the addition of 10 μL 2.5 mM NADPH into each well. Before the addition of NADPH, the fluorescence intensity of each well (t = 0) was measured with the Tecan Infinite M200 Microplate Reader (Tecan Group Ltd, Mannedorf, Switzerland) using 380 nm excitation and 455 nm emission filters, and was considered as basal fluorescence. The microplates were then incubated at room temperature (22 ± 1°C) and the fluorescence intensity was determined after 1 h. The reaction was stopped by adding 100 μL 50% (v/v) glycine buffer (0.5 M)-ethanol, pH 10.4 and fluorescence (t = 1 h). Relative fluorescence units (RFU) was calculated as RFU = fluorescence (t = 1 h) - fluorescence (t = 0). Three or four replicates were performed for each treatment for all assays in this study.

### Optimization of the P450 activity assay conditions

We found that homogenizing *M. persicae* in the microplate resulted in significantly higher RFU; therefore, the following assays were conducted to optimize the microplate homogenization method. First, we tested the necessity of NADPH and protectants (i.e. EDTA, DTT, PMSF, and glycerol) in P450 activity measurement. Three adult *M. persicae* were homogenized in each well of the microplate using pipette tips in 50 μL 100 mM phosphate buffer or with phosphate buffer containing protectants. Then 50 μL 7-EC solutions (1 mM 7-EC dissolved in 100 mM phosphate buffer pH 7.5 with or without protectants) were added to each well. The reaction was initiated by the addition of 10 μL 2.5 mM NADPH or 10 μL phosphate buffer and stopped by adding 100 μL 50% (v/v) glycine buffer (0.5 M)-ethanol after 1 h. The RFU was determined by the microplate reader as described above.

Because we found that NADPH and protectants was not indispensable in measuring aphid P450 activity, these chemicals were not used in the following experiments. To optimize the homogenization method, three adult *M. persicae* were placed in each microplate well and were gently crushed or ground with pipette tips for different cycles. To optimize the number of aphids used in each well, 1, 2, 3, or 5 adult *M. persicae* were homogenized in the microplate and their P450 activities were determined as described above. Time dependence of *M. persicae* P450 activities were determined similarly by using two adult aphids in each microplate well.

### Inhibitory effect of *M. persicae* homogenate

Two mg aphid equivalents homogenized in the EP tube without centrifugation were mixed with liver extracts to examine whether the lower P450 activity of aphids homogenized in the EP tube was due to inhibitors released during homogenization. Liver extract was prepared by grinding fresh pig liver tissue in 100 mM phosphate buffer (pH 7.5) using a glass mortar and pestle. The liver homogenate was then centrifuged at 4°C 13,000 g for 15 min and the supernatant was collected and used. To test the heat stability of the endogenous inhibitors, aphid homogenate was heated in boiling water for 5 min and then mixed with liver extract. The RFU was determined and calculated following previous procedure using 7-EC (dissolved in 100 mM phosphate buffer pH 7.5) as substrate.

We also tested the inhibitory effect of the supernatants of aphid homogenate and whether homogenizing aphids in the microplate had such inhibitory effect on liver extract. After being homogenized in the EP tube, aphid homogenate was centrifuged at 4°C 13,000 g for 15 min and the supernatant was collected and used. For microplate homogenization method, aphids were ground in the microplate using pipette tips for 20 cycles and then mixed with liver extract. The RFU was determined and calculated as described above.

### Suitability of the microplate homogenization method for *A. pisum* and *R. padi*

To test the suitability and necessity of the microplate homogenization method for measuring P450 activities in *A. pisum* and *R. padi*, ECOD activities in these aphids were determined and compared using different homogenization methods. The inhibitory effects of both aphid homogenates on the P450 activity of liver extract were also examined.

### Fluorescence of chemicals involved in the P450 activity assay

One hundred μL 7-EC, 7-OH, or NADPH, dissolved in 100 mM phosphate buffer pH 7.5 with different concentrations, were put into the microplate and the fluorescence was determined with the microplate reader. The phosphate buffer was used as the blank control. The fluorescence of these chemicals was also measured using similar methods after the addition of 100 μL 50% (v/v) glycine buffer (0.5 M)-ethanol, pH 10.4.

### Statistical analysis

The RFU of *M. persicae*, *A. pisum*, and *R. padi* homogenized with different methods were analyzed using Student’s t-test. The influence of NADPH and protectants, grinding degree, number of aphids used in each microplate well, and inhibitory effects of aphid homogenate on ECOD activities were analyzed using one-way analysis of variance (ANOVA), followed by Fisher’s least significant difference (LSD) tests, at a significance level of P < 0.05. All statistical analyses were performed using the IBM SPSS Statistics package 19 (SPSS Inc., Chicago, IL, USA).

## Results and discussion

### Comparison of homogenization methods

The ECOD activity in *M. persicae* was about three times higher when homogenized aphids in the microplate than in the EP tube (Figure 1; df = 4; t = 8.28; P = 0.001), which agrees with a previous study that failed to detect ECOD activities in *M. persicae* because of endogenous inhibitors released during homogenization (Philippou et al., 2010). In addition, we found that homogenized *M. persicae* in the EP tube (without centrifugation) reduced liver ECOD activities by 90% (Figure 2A; df = 2, 9; F = 427.806; P < 0.001), and this effect still existed after heating aphid homogenate in boiling water (Figure 2A) or after centrifugation (Figure 2B). According to previous studies, the inhibitory compounds in *M. persicae* could be eye pigment and/or uncharacterized nucleoprotein (Wilson & Hodgson, 1972; Gilbert & Wilkinson, 1975). However, aphids homogenized in the 96-well microplate showed higher ECOD activities (Figures 1), and did not inhibit liver ECOD activities (Figure 2B). Homogenizing aphids in the microplate may not completely grind aphid tissues and thus likely release little or no endogenous inhibitors. These findings suggest the importance of considering endogenous inhibitors before measuring cytochrome P450 activity in aphids.

**Figure 1.**
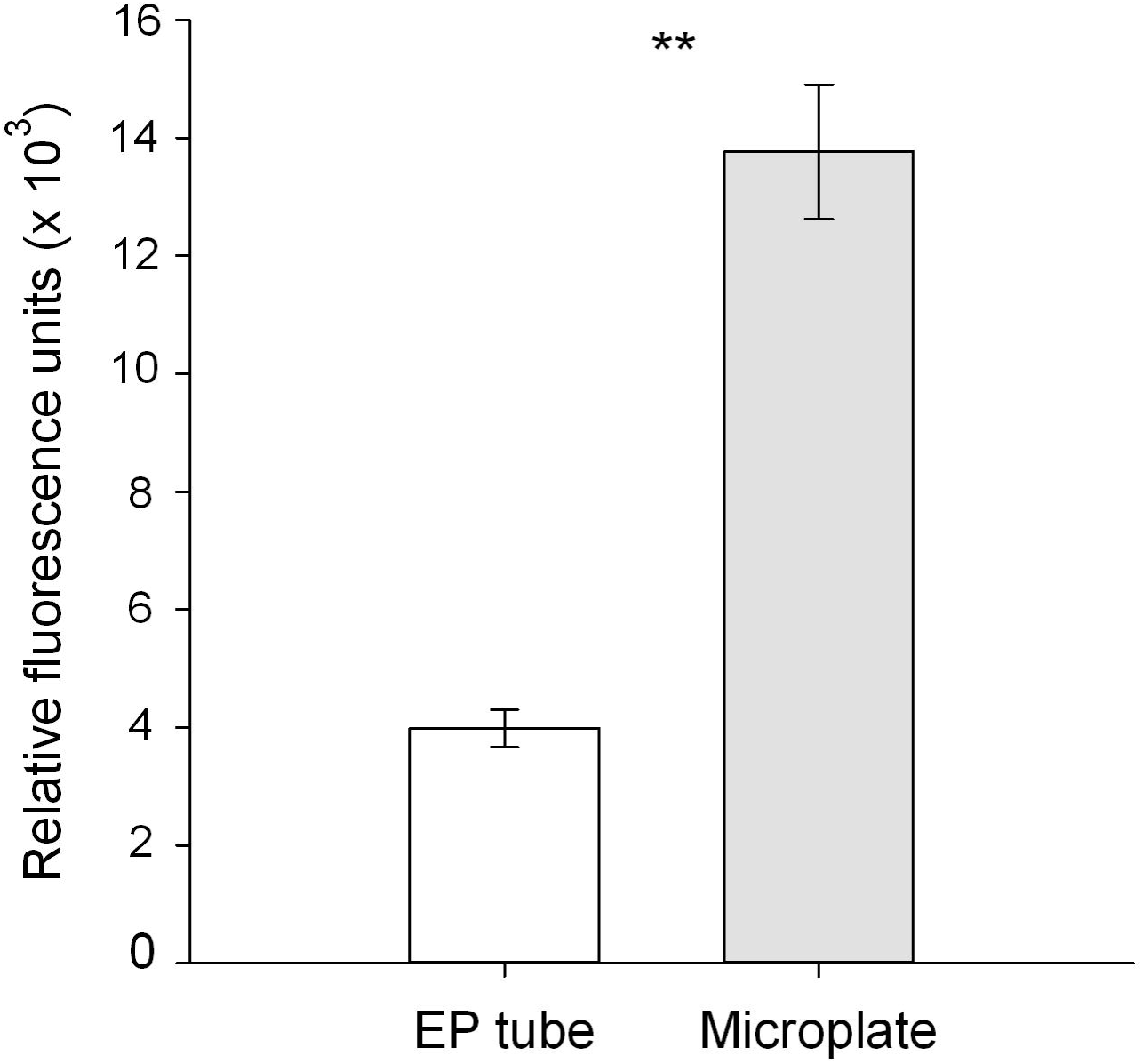
ECOD activities of *M. persicae* homogenized with different methods. Aphids were homogenized in the EP tube with fitted pestles or ground in 96-well microplate using pipette tips. Data are mean ± SE (n = 3). ^∗∗^P < 0.01 (Student’s t-test).

**Figure 2.**
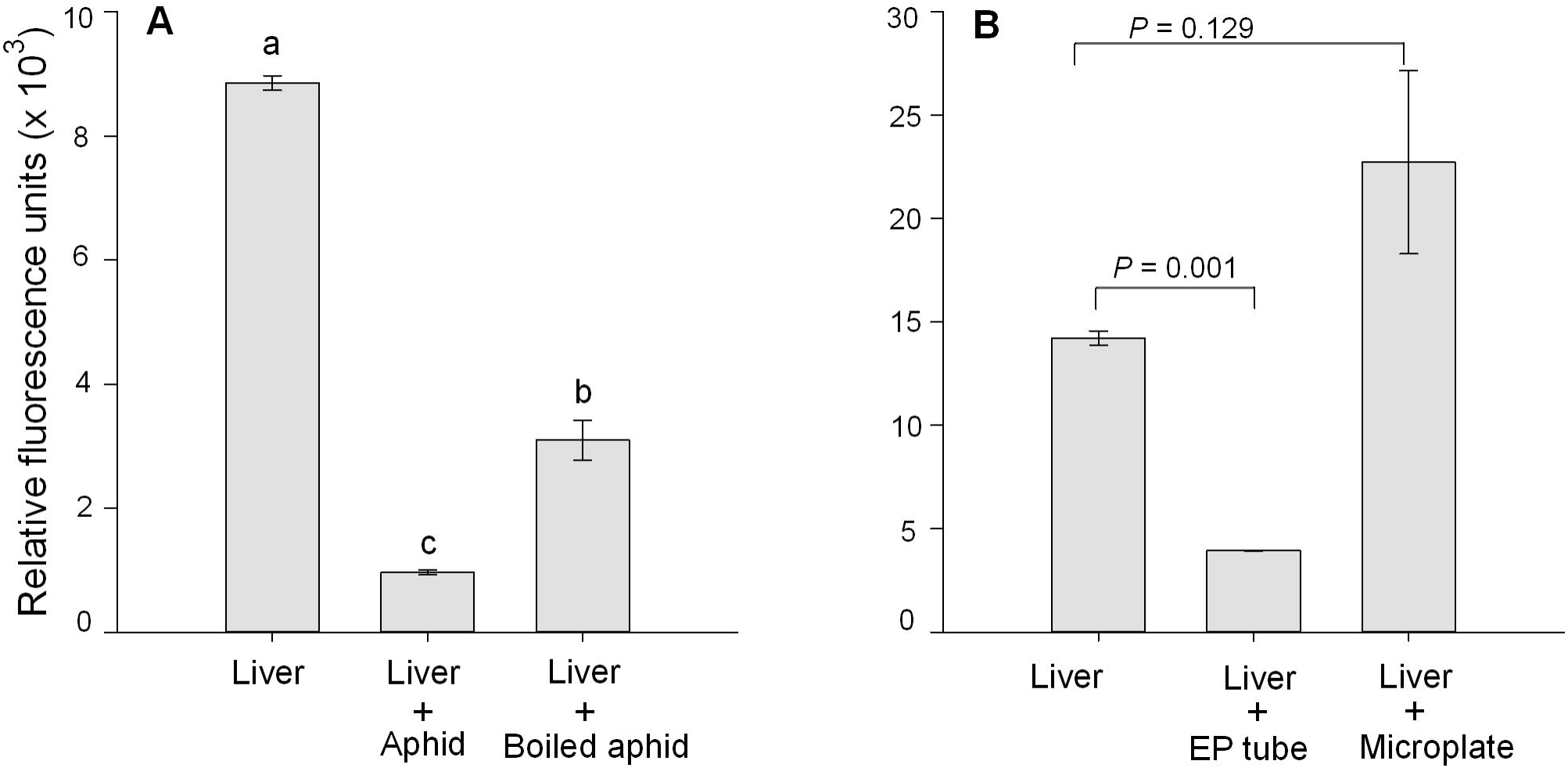
Inhibitory effects of *M. persicae* homogenate on ECOD activity of liver extract. (A) Aphids homogenized in the EP tube (without centrifugation) reduced liver ECOD activity (Liver + Aphid) and the inhibitory effects still existed after heating aphid homogenate in boiling water (Liver + Boiled aphid). (B) Aphids homogenized in the EP tube (after centrifugation) reduced liver ECOD activities (Liver + EP tube), whereas aphids that were directly homogenized in the microplate had no such inhibitory effect (Liver + Microplate). Data are mean ± SE (n = 4). Different letters above bars indicate significant difference at P < 0.05 (one-way ANOVA followed by LSD test). The P values above bars was calculated by Student’s t-test.

### Optimization of the ECOD activity assay protocol

Since aphid ECOD activities were not enhanced after the addition of the protectants and NADPH, these chemicals are not indispensable in measuring aphid ECOD activity (Figure 3A; df = 2, 6; F = 3.945; P = 0.081). Aphid tissues homogenized in the microplate possibly generated enough NADPH for the P450 activity assay and thus NADPH was not needed (Desousa et al., 1995; Castaneda et al., 2009). Moreover, the fluorescence of residue NADPH may affect the accuracy of the P450 activity assay and needs further processes to eliminate this influence (Chauret et al., 1999).

**Figure 3.**
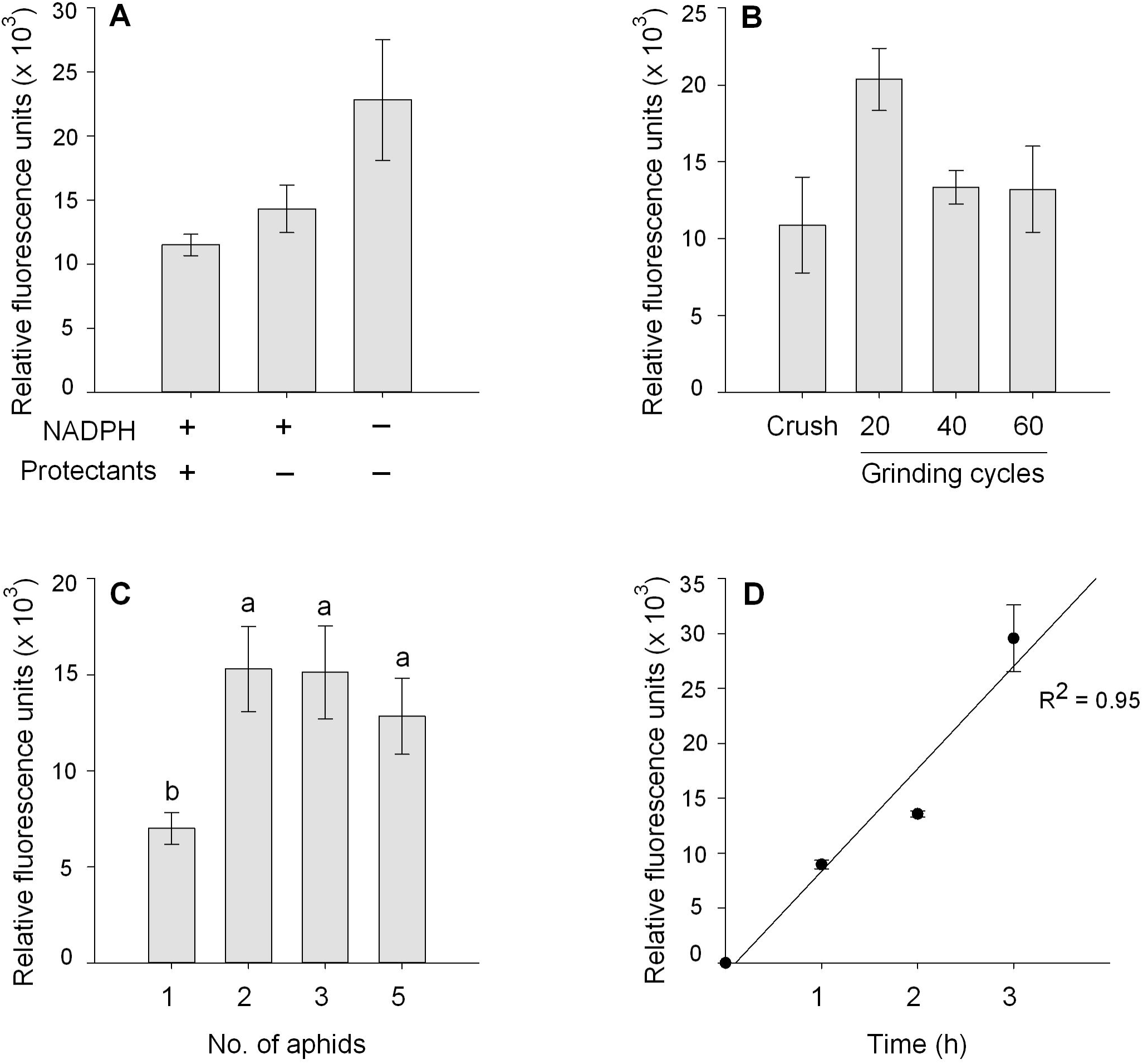
Optimization of the P450 enzyme assay conditions. The influence of (A) NADPH and protectants, (B) grinding degree, (C) number of aphids in each microplate well, and (D) reaction time on ECOD activities of *M. persicae.* Data are mean ± SE (n = 3). Different letters above bars indicate significant difference at P < 0.05 (one-way ANOVA followed by LSD test).

Grinding *M. persicae* for 20 cycles in the microplate resulted in relatively higher ECOD activities, whereas higher grinding cycles or crush alone did not lead to higher ECOD activities (Figure 3B; df = 3, 8; F = 2.943; P = 0.099). Homogenizing aphids by crushing alone may limit the contact of aphid P450 enzyme with 7-EC, while more grinding cycles may release more inhibitors, both of which lead to lower ECOD activities. Similarly, three or five aphids in one microplate well did not generate higher ECOD activities than two aphids (Figure 3C; df = 3, 15; F = 5.949; P = 0.007), which may be due to that more aphids in one microplate well lead to higher concentration of inhibitors. Although the RFU of this assay remained linear for 3 h (Figure 3D; R^2^ = 0.95), the ECOD activities of different samples could be differentiated in 1 h because of the high sensitivity. In summary, we improved the method measuring aphid P450 enzyme activities as follows: one or two aphids were placed in one well of a 96-well microplate, and ground in phosphate buffer using pipette tips for 20 cycles, followed by addition of 7-EC, and then incubated for 1 h at room temperature before adding glycine buffer-ethanol mixture to stop the reaction.

### Suitability of the microplate homogenization method for *A. pisum* and *R. padi*

The pea aphid, *A. pisum* had only approximately one third ECOD activities when homogenized in the EP tube than in the microplate (Figure 4A; df = 6; t = 3.208; P = 0.018), and this homogenate significantly reduced liver ECOD activities (Figure 4C), suggesting that *A. pisum* released P450 enzyme inhibitors during homogenization in the EP tube. However, the bird cherry-oat aphid, *R. padi*, had no such inhibitory effect on liver ECOD activities (Figure 4C) and showed comparable ECOD activities (Figure 4B; df = 6; t = 0.079; P = 0.939) when homogenized with different methods, implying that there may be few or no P450 enzyme inhibitors in *R. padi.*

**Figure 4.**
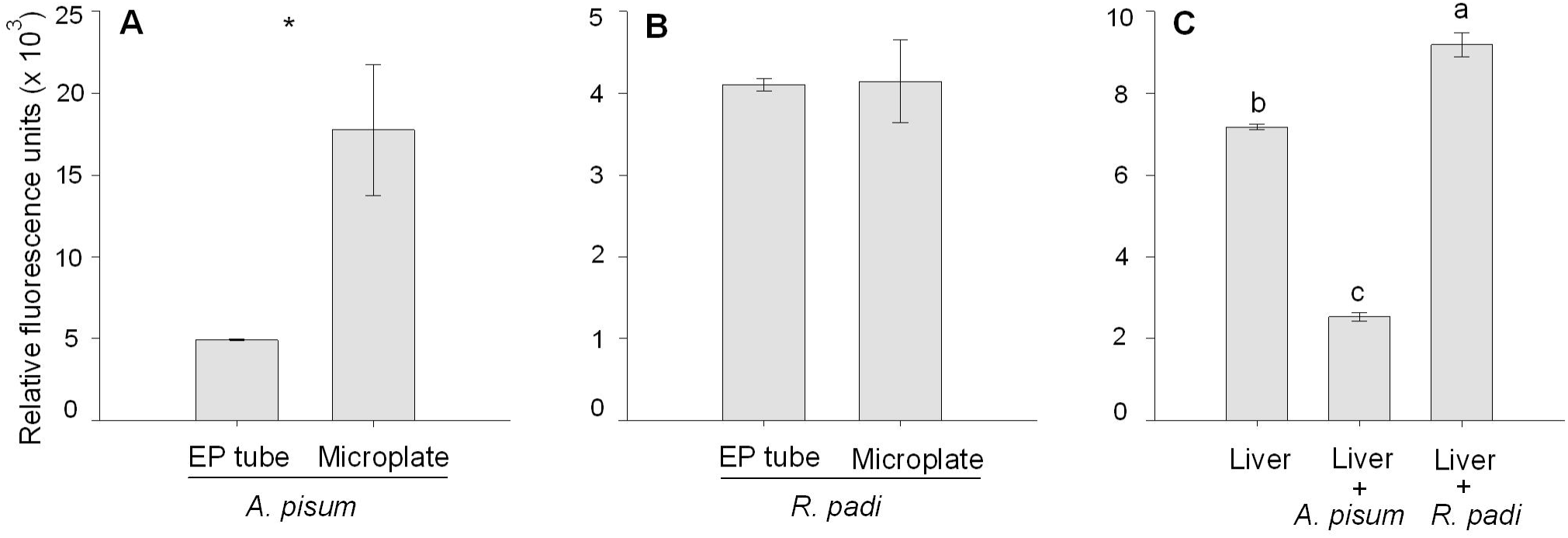
ECOD activities of (A) *A. pisum* and (B) *R. padi* homogenized in the EP tube or microplate and (C) inhibitory effects of aphid homogenate (homogenized in the EP tube) on liver ECOD activity. Data are mean ± SE (n = 4). ^∗^P < 0.05 (Student’s t-test) and different letters above bars indicate significant difference at P < 0.05 (one-way ANOVA followed by LSD test).

### Fluorescence of chemicals involved in the P450 activity assay

The P450 enzyme substrate 7-EC had a low fluorescence intensity (Figure 5A; 1.27 a.u./μM), and its fluorescence intensity increased slightly after the addition of glycine buffer-ethanol mixture (Figure 5A; 1.69 a.u./μM). However, 7-OH had a relative higher fluorescence levels (Figure 5B; 12,083 a.u./μM) and the fluorescence intensity increased 3.7 times after the addition of glycine buffer-ethanol mixture (Figure 5B; 44,645 a.u./μM). These results suggest that 7-EC is suitable as the P450 enzyme substrate, and the high pH glycine buffer-ethanol buffer can improve the sensitivity of such assay. The NADPH should not be used in this assay, because NADPH is not needed and had a relative high fluorescence intensity (Figure 5C; 9.72 a.u./μM).

**Figure 5.**
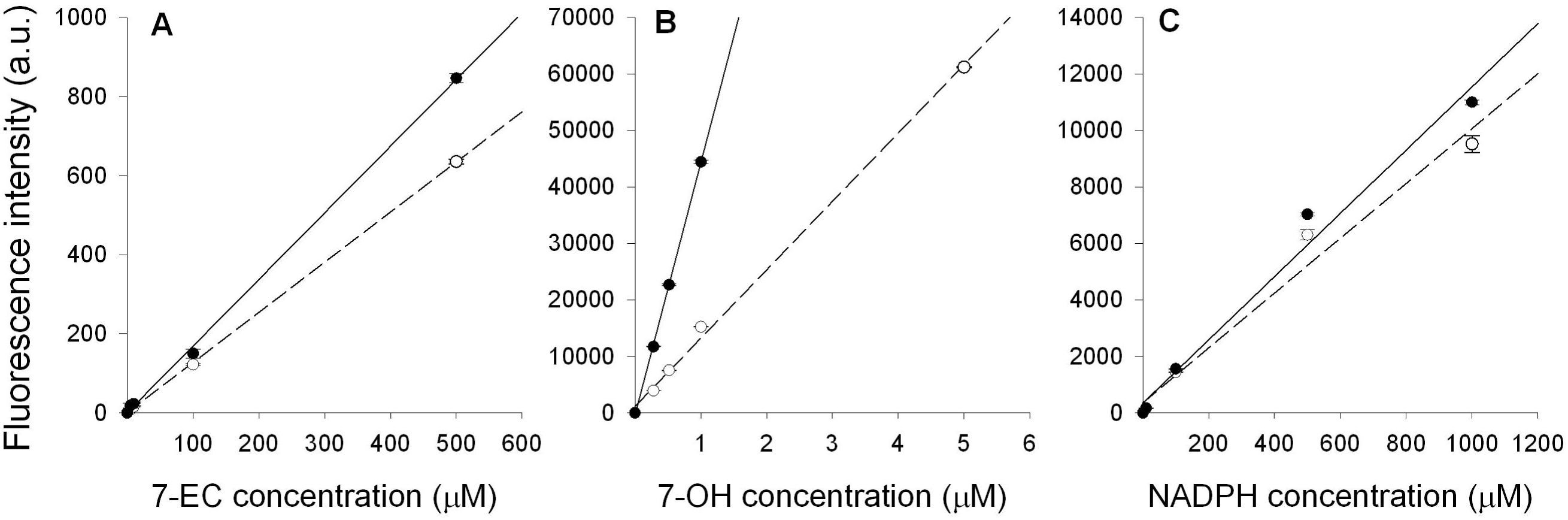
Fluorescence intensity of (A)7-EC, (B) 7-OH, and (C) NADPH before and after the addition of 50% (v/v) glycine buffer-ethanol. Data are mean ± SE (n = 3).

## Conclusion

Although homogenizing aphids in the 96-well microplate reduced the release of endogenous inhibitors, it is difficult to guarantee that no inhibitors were released. However, the samples containing two aphids had about two times higher ECOD activities than one aphid, suggesting that our method can eliminate most, if not all, inhibitors. Nevertheless, the number of grinding cycles and number of aphids in each microplate well should be optimized before specific P450 activity assay. This study suggests the necessary of testing the inhibitory effects of endogenous compounds before measuring P450 activity, and provides one possible method to reduce this inhibitory effect. We used aphids in the present study, but our method may be also suitable for other organisms in the P450 activity assay.

## Acknowledgments

Funding of this research was supported by the China Postdoctoral Science Foundation Grant (No. 2017M613229). We are grateful for the assistance of all staff and students in the Key Laboratory of Applied Entomology, Northwest A&F University at Yangling, Shaanxi, China.

## References

Castaneda LE, Figueroa CC, Fuentes-Contreras E, Niemeyer HM & Nespolo RF (2009) Energetic costs of detoxification systems in herbivores feeding on chemically defended host plants: a correlational study in the grain aphid, *Sitobion avenae*. Journal of Experimental Biology 212: 1185–1190.

Chauret N, Tremblay N, Lackman RL, Gauthier J-Y, Silva JM et al. (1999) Description of a 96-well plate assay to measure cytochrome P4503a inhibition in human liver microsomes using a selective fluorescent probe. Analytical Biochemistry 276: 215–226.

Desousa G, Cuany A, Brun A, Amichot M, Rahmani R & Bergé J-B (1995) A microfluorometric method for measuring ethoxycoumarin-O-deethylase activity on individual *Drosophila melanogaster* abdomens: interest for screening resistance in insect populations. Analytical Biochemistry 229: 86–91.

Gilbert MD & Wilkinson CF (1975) An inhibitor of microsomal oxidation from gut tissues of the honey bee (*Apis mellifera*). Comparative Biochemistry and Physiology Part B: Comparative Biochemistry 50: 613–619.

Gottardi M, Kretschmann A & Cedergreen N (2016) Measuring cytochrome P450 activity in aquatic invertebrates: a critical evaluation of in vitro and in vivo methods. Ecotoxicology 25: 419–430.

Liu N, Li M, Gong Y, Liu F & Li T (2015) Cytochrome P450s-Their expression, regulation, and role in insecticide resistance. Pesticide Biochemistry and Physiology 120: 77–81.

Orrenius S, Berggren M, Moldéus P & Krieger RI (1971) Mechanism of inhibition of microsomal mixed-function oxidation by the gut-contents inhibitor of the southern armyworm (*Prodenia eridania*). Biochemical Journal 124: 427–430.

Pasay C, Arlian L, Morgan M, Gunning R, Rossiter L et al. (2009) The effect of insecticide synergists on the response of scabies mites to pyrethroid acaricides. PLoS Neglected Tropical Diseases 3: e354. doi:10.1371/journal.pntd.0000354.

Philippou D, Field L & Moores G (2010) Metabolic enzyme(s) confer imidacloprid resistance in a clone of *Myzus persicae* (Sulzer) (Hemiptera: Aphididae) from Greece. Pest Management Science 66: 390–395.

Puinean AM, Foster SP, Oliphant L, Denholm I, Field LM et al. (2010) Amplification of a cytochrome P450 gene is associated with resistance to neonicotinoid insecticides in the aphid *Myzus persicae*. PloS Genetics 6. doi:10.1371/journal.pgen.1000999.

Valles SM & Yu SJ (1996) German cockroach (Dictyoptera: Blattellidae) gut contents inhibit cytochrome P450 monooxygenases. Journal of Economic Entomology 89: 1508–1512.

Wilson TG & Hodgson E (1972) Mechanism of microsomal mixed-function oxidase inhibitor from the housefly *Musca domestica* L. Pesticide Biochemistry and Physiology 2: 64–71.

